# A dynamic model of neurovascular coupling network accounting for capillary *K*^+^-sensing vasomotion

**DOI:** 10.1101/2023.08.04.551924

**Authors:** Zhixuan Yuan, Yangyang Yu, Ying Wu

**Affiliations:** State Key Laboratory for Strength and Vibration of Mechanical Structures Shaanxi Engineering Laboratory for Vibration Control of Aerospace Structures School of Aerospace Engineering, Xi’an Jiaotong University, Xi’an 710049, China

## Abstract

Neurovascular coupling (NVC) is a fundamental unit that elucidates the regulation of vascular activity by neuronal electrical activity, ensuring adequate blood flow to the brain’s active regions and optimizing neural function. In this article, we propose a comprehensive dynamic model of a neural vascular coupling network that integrates the capillary *K*^+^ sensing mechanism and its impact on vascular motion. Our computational framework incorporates the dynamics of neurons, astrocytes, and smooth muscle cells at the cellular level. By integrating recent experimental findings, we have successfully demonstrated the ability of capillary endothelial cells to detect localized fluctuations in extracellular potassium (*K*^+^) concentration via the Kir2.1 channel situated on their membranes. Subsequently, these cells transmit this signal to smooth muscle cells, initiating the release of vasoactive substances and thereby facilitating vascular movement. This sequence of events serves as the fundamental basis of our model. Furthermore, the simulation we provide supports the inclusion of a temperature term in our model, enabling accurate replication of experimental observations: variations in neuronal electrical activity resulting from an increase in temperature and subsequent vasoconstriction. In conclusion, our dynamic model presents a valuable tool for investigating the mechanisms underlying neurovascular coupling and the role of capillary *K*^+^ sensing in regulating vascular activity. It enhances our understanding of brain function and offers insights for developing therapeutic interventions for neurovascular diseases.

**Author summary:** It is widely accepted that neuronal activity induces vasodilation, resulting in increased blood flow, while vascular activity provides energy supply to the nervous system. The role of astrocytes in these processes has been extensively studied, revealing their dual involvement in neuronal firing and facilitation of energy delivery to the nervous system through blood vessels. As cellular mediators of bidirectional processes, astrocytes play a crucial role in promoting neurovascular coupling, leading to the formation of neurovascular coupling units. Building upon existing experimental observations and previous theoretical reports, the present study successfully replicates the phenomenon of vasodilation caused by an increase in potassium ion concentration in the perivascular space, as well as vasoconstriction resulting from elevated potassium ion concentration. A new neuron-astrocyte-endothelial-smooth muscle cell coupling network model, incorporating temperature considerations, was constructed to achieve these outcomes. Additionally, by manipulating temperature, the experimental observation of neuronal vasoconstriction during febrile seizures was successfully reproduced. These results highlight the significance of the neurovascular coupling unit as the fundamental entity for investigating the functions and mechanisms of the nervous system and brain.

## Introduction

Neurovascular coupling (NVC) is the process by which neuronal activity regulates blood flow in the brain, ensuring that adequate oxygen and nutrients are delivered to active regions [1–4]. This process is essential for normal brain function, and disturbances in NVC have been implicated in various neurological disorders [5–8]. NVC is mediated by a complex network of cells, including neurons, astrocytes, pericytes, endothelial cells (ECs), and smooth muscle cells (SMCs), which communicate through multiple signaling pathways [1, 9].

Capillary *K*^+^-sensing vasomotion is a recently discovered mechanism by which capillary ECs detect and respond to changes in perivascular potassium (*K*^+^) levels through inward rectifier potassium (Kir) channels [10–12]. The subsequent membrane potential depolarization in ECs is transmitted to adjacent smooth muscle cells (SMCs) through gap junctions at myoendothelial projections [13–15]. This depolarization causes SMCs to become hyperpolarized, causing the inactivation of voltage-dependent calcium (*Ca*^2+^) channels. As a result, there is a decrease in *Ca*^2+^ influx and a reduction in *Ca*^2+^-dependent actin-myosin cross-bridge cycling, leading to vasodilation [16].

Potassium (*K*^+^) channels play a critical role in regulating the membrane potential of various types of cells [17, 18], including smooth muscle and endothelial cells [19]. Among the different types of *K*^+^ channels, the inwardly rectifying *K*^+^ (Kir) channels have been shown to be particularly important in regulating the electrical activity, contractility of smooth muscle cells and endothelial cells, and the release of vasoactive substances [20–22]. Kir channels exhibit inward rectification, whereby they conduct *K*^+^ ions more efficiently in the inward direction (toward the cell interior) than in the outward direction, resulting in an inward rectification of the current-voltage relationship. Within the Kir channel family, Kir2.1 channels in endothelial cells (ECs) rather than smooth muscle cells (SMCs) are critical in regulating vascular tone and blood pressure in the brain [23], especially during periods of increased neuronal activity [24].

There are numerous types of epilepsy, and over recent years, research on epilepsy modeling has witnessed a steady increase [25–29]. Febrile status epilepticus (FSE) constitutes a significant risk factor for the development of temporal lobe epilepsy. While the majority of febrile seizures are typically transient and devoid of long-term consequences, seizures lasting beyond 30 minutes fall under the classification of febrile status epilepticus (FSE) [30, 31]. The temperature term model and the initial HH model were already established in previous studies [32]. Yu used a temperature model to demonstrate that the increase in energy efficiency is primarily attributed to elevated body temperature [33]. In a separate study, Du discovered that incorporating a temperature term into the channel model revealed the temperature-dependent dysfunction of the astrocyte Kir4.1 channel, which may contribute to febrile seizures [34]. The temperature term utilized in this article has been refined and modified based on the methodologies described in the two aforementioned studies.

Dynamic modeling of neurovascular coupling system has gained significant attention in recent years [35–37]. Chander established a computational model of the neuro-glio-vascular system, which assumed that the change in vascular radius was only linearly related to the membrane potential of smooth muscle cells [38]. Bennett constructed a metabolic neurovascular coupling unit in which the release of vasoactive factors occurred outside the astrocytes rather than inside [39]. However, the vasomotor mechanisms induced by extravascular high potassium ions have not been fully elucidated. In this paper, we propose a novel dynamic model of a neurovascular coupling network that takes into account the influence of temperature, integrating the motion of blood vessels triggered by capillary *K*^+^. Our model includes multiple cell types, such as neurons, astrocytes, endothelial cells, and smooth muscle cells, as well as the extracellular spaces, and focuses on Kir2.1 in endothelial cells.

Utilizing the aforementioned NVC model, we conducted a simulation investigating the vascular response resulting from alterations in perivascular potassium ion concentration within the experimental setup [15, 40]. Furthermore, upon taking into account the influence of temperature, our simulation outcomes exhibited a high degree of concordance with the vasoconstrictive effects induced by the thermal convulsion as delineated in the experimental observations [41]. Overall, our study offers new insights into the mechanisms underlying neurovascular coupling and emphasizes the significance of capillary *K*^+^-sensing vasomotion in regulating blood flow in the brain. Furthermore, our model may serve as a valuable tool for investigating the role of NVC in neurological disorders and developing new therapeutic strategies.

## Results

### Schematic diagram of the model

The entire neurovascular coupling (NVC) model is comprised of six components: a neuron, the neuronal extracellular space (ECS), an astrocyte, the perivascular space (PVS), an endothelial cell (EC), and a smooth muscle cell (SMC) (Fig 1). As shown in Fig 1, neuronal electrical activity induces the exchange of ions and the release of the neurotransmitter glutamate (Glu). The astrocyte absorbs excess *K*^+^ via Kir4.1 channels, while extracellular Glu activates mGluR5 on the membrane, resulting in the outflow of *Ca*^2+^ from the calcium store ER. Elevated *Ca*^2+^ levels, together with their produced epoxyeicosatrienoic acid (EET), activate the BK channel in astrocytic endfeet. *K*^+^ in the PVS promotes the activation of the Kir2.1 channel on the EC and the VGPC on the SMC. Subsequently, the hyperpolarization of the EC membrane generated by the Kir2.1 channel transfers to the SMC via myoendothelial gap junctions [14], causing a decrease in intracellular *Ca*^2+^. The calcium-influenced actin-myosin cross-bridge cycle ultimately results in vasomotion.

**Fig 1.**
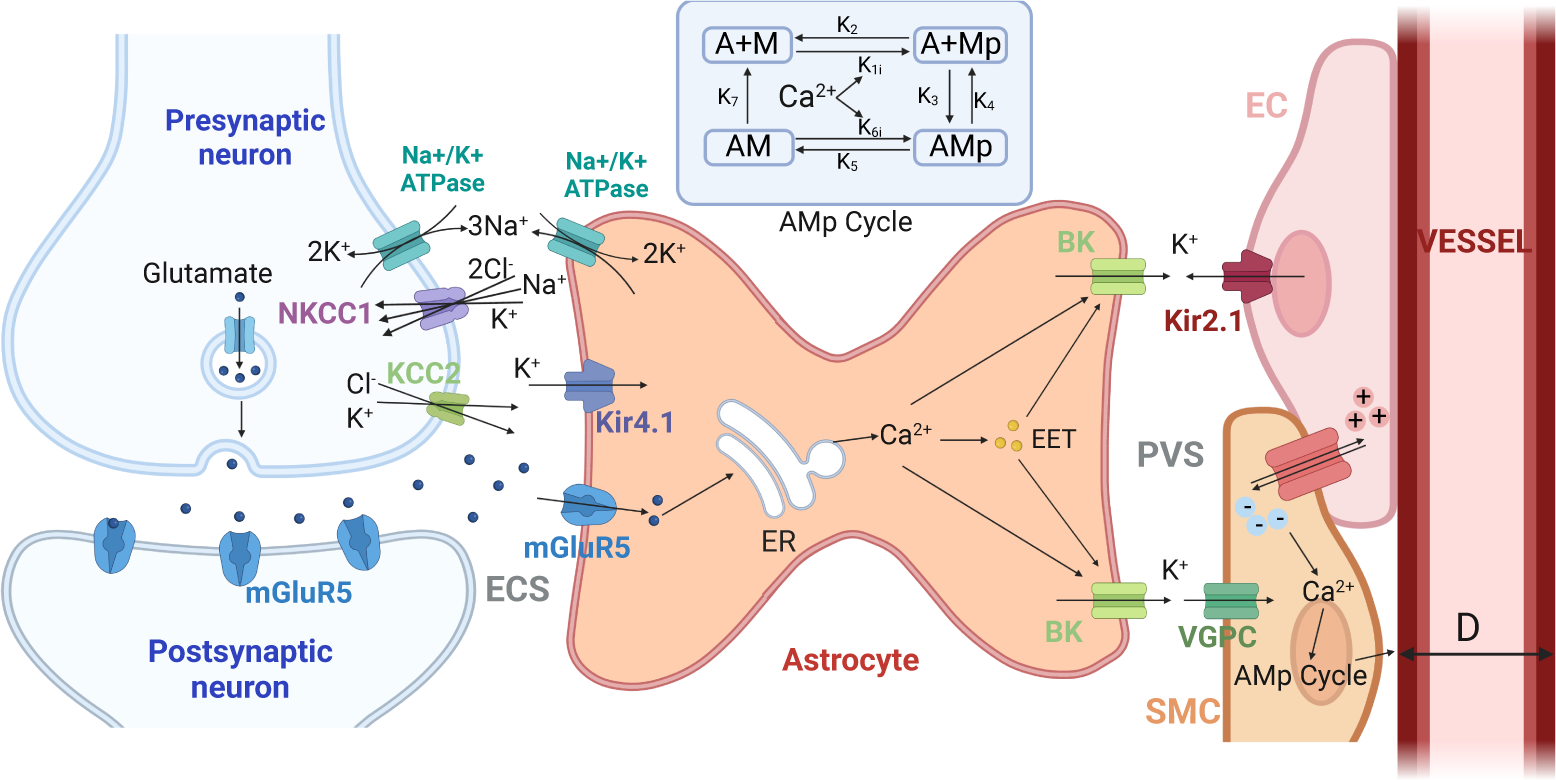
The diagram of this neurovascular coupling (NVC) model including a neuron, the extracellular space (ECS), an astrocyte, the perivascular space (PVS), an endothelial cell (EC) and a smooth muscle cell (SMC). NKCC1: *Na*^+^/*K*^+^/*Cl^−^* cotransporter, the transportation ratio is 1*Na*^+^:1*K*^+^:2*Cl^−^*; KCC2: *K*^+^/*Cl^−^* cotransporter, the transportation ratio is 1*K*^+^:1*Cl^−^*; Kir4.1 and Kir2.1 channels: inward rectifier potassium ion channels; mGluR5: metabotropic glutamate receptor 5, a type of G protein-coupled neurotransmitter glutamate receptor; ER: endoplasmic reticulum; EET: epoxyeicosatrienoic acid, a kind of vasoactive factor; BK: large conductance calcium-activated potassium channel; VGPC: voltage-gated potassium channel. Created by Biorender.com.

### Modelling Kir2.1 on endothelial cells

At present, no models of Kir2.1 channels on ECs exist. However, we started by building on the Kir4.1 channel model established for astrocytes [42]. Using this as a foundation, we obtained a basic Kir2.1 model with inward rectification properties:

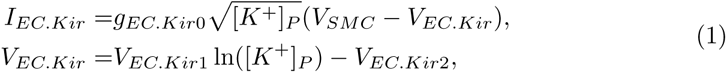

The basic model represents the Kir2.1 channel affected by the *K*^+^ in the perivascular space. Experimental recordings have shown that changes in perivascular *K*^+^ concentration cause vasomotion (see Fig 2(a)), we observed that compared to normal physiological conditions([*K*^+^]*_P_* =6mM), elevated *K*^+^ concentration can lead to vasodilation([*K*^+^]*_P_* =16mM), but excessive *K*^+^ concentration can inhibit this dilation effect([*K*^+^]*_P_* =21mM), and even lead to vasoconstriction([*K*^+^]*_P_* =26mM). Given the double-edged sword characteristics of *K*^+^ outside, we assume that the Kir2.1 channel has dual gating:

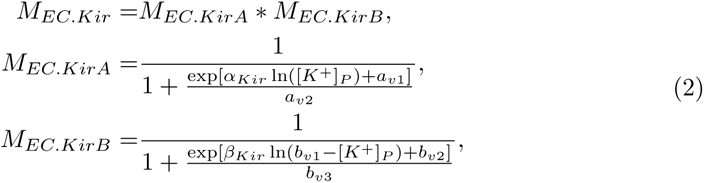

**Fig 2.**
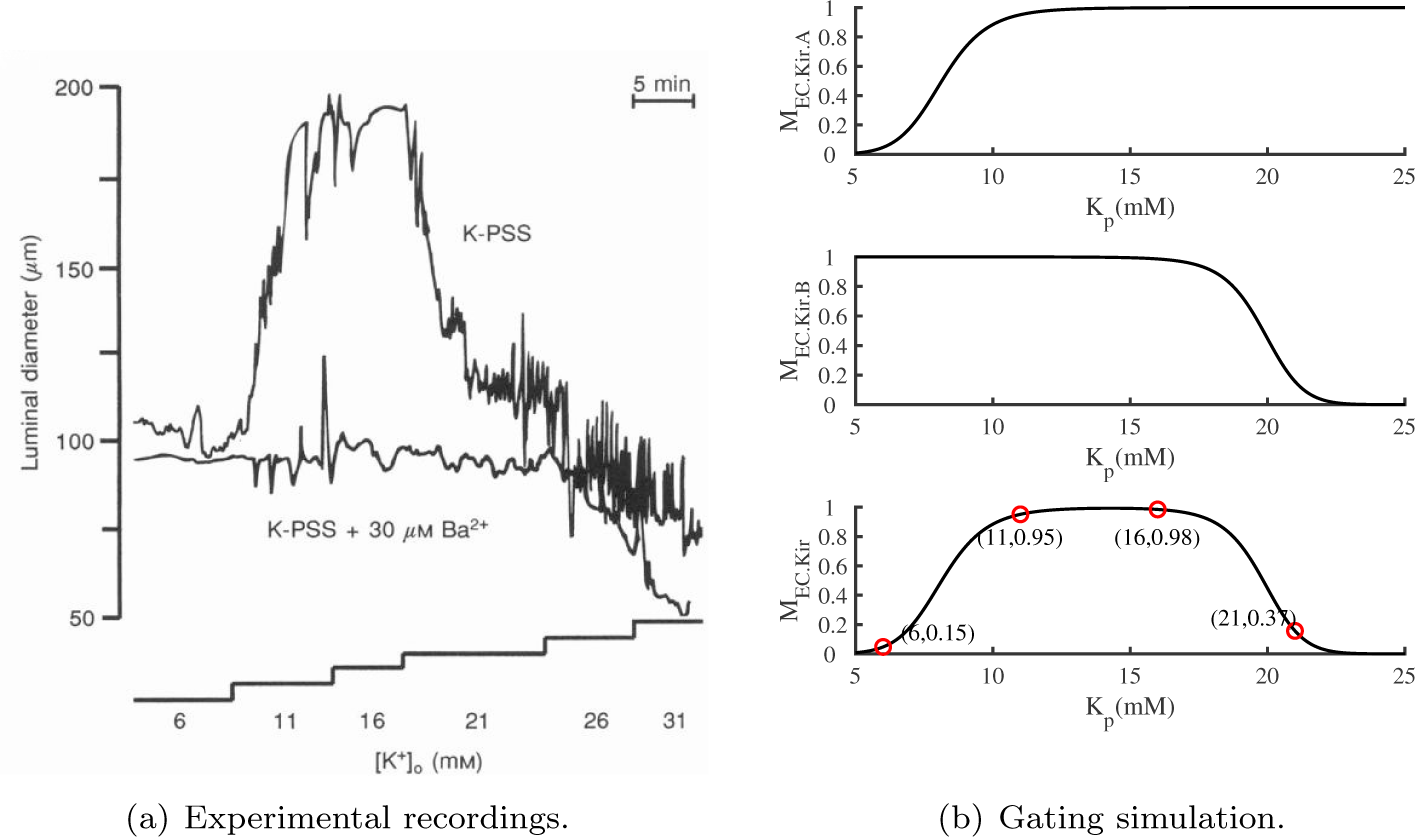
(a) Fig 3(c) from Ref. [40] illustrates the effect of graded increases of [*K*^+^]*_o_* (referred to as [*K*^+^]*_P_* in this paper) on the diameter of a cerebral artery in K-PSS(*K*^+^ physiological salt solution) (upper trace). The lower trace shows the response of this artery to increases in [*K*^+^]*_o_* in the presence of 30*µ*M *Ba*^2+^ (Kir channel inhibitor) added to the bathing solution. (b) The steady-state opening of the gates M in our model varies with [*K*^+^]*_P_*, and the corresponding gate values for specific [*K*^+^]*_P_* are marked with red circles.

The gating diagram, obtained by continuously selecting and debugging the parameters in Eqn.(2), is shown in Fig 2(b), along with the gating values that vary with potassium ions. Thus far, we have derived a Kir2.1 channel model that exhibits both inward rectification for potassium ions and dual gating behavior:

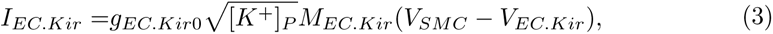

Next, we will demonstrate the validity of the model by focusing on its qualitative aspects. By varying the parameter [*K*^+^]*_P_*_0_ in Eqn.(21) (see S1 Appendix) to different values, the *K*^+^ concentration in the perivascular space is adjusted to match that of Fig 2, namely, approximately 6, 11, 16, 21, and 26 *mM*.

The results, depicted in the Fig 3, demonstrate that with the increase in the value of [*K*^+^]*_P_*, blood vessels experience dilation, suppressed dilation and contraction. Fig 3(d) exhibits a larger span of change in blood vessel diameter compared to the other four figures. This is attributed to the fluctuation in [*K*^+^]*_P_* concentration occurring within the range where the gate value undergoes significant changes. The diameter of the blood vessel in Fig 3(e) is also reduced compared to that in Fig 3(a) because of the complete closure of the Kir2.1 channel in (e), rather than in (a). To facilitate clearer observation, we have represented the potassium ion concentration and blood vessel diameter as the horizontal and vertical axes, respectively, in Fig 4, as shown in Fig 3.

**Fig 3.**
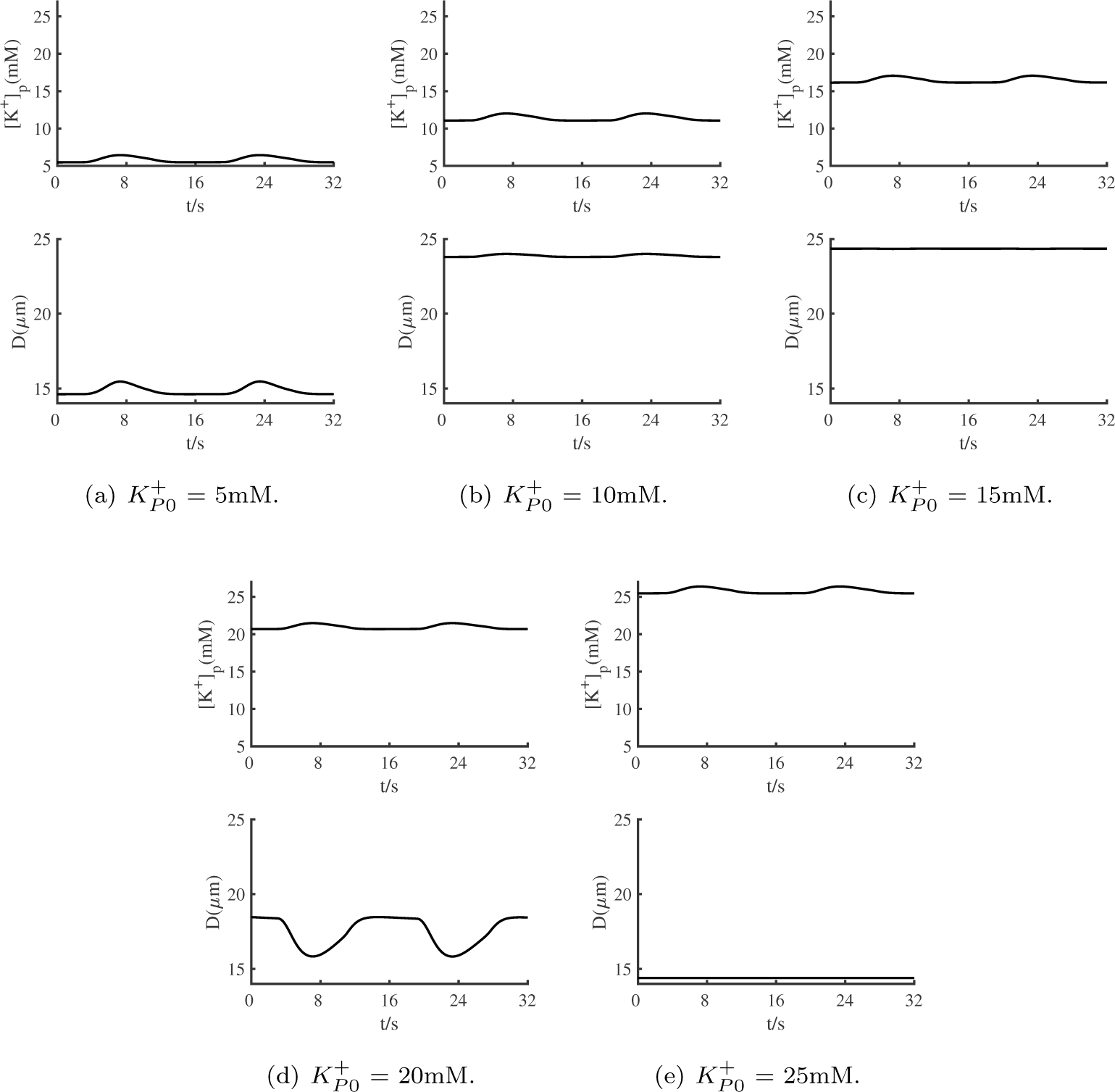
Responses of perivascular potassium ion concentration ([*K*^+^]*_P_*) and vessel diameter (*D*) induced by different [*K*^+^]*_P_*_0_.

**Fig 4.**
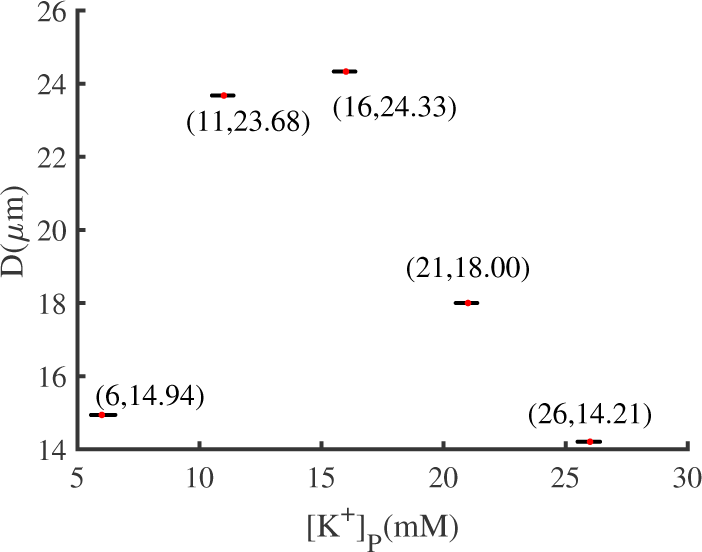
Potassium concentration in the perivascular space versus vessel diameter in Fig 3(a)-(e).

Fig 4, constructed from Fig 3, clearly illustrates that an increase in [*K*^+^]*_P_* concentration leads to a expansion followed by constriction of the blood vessel diameter. Simulation results in Fig 4 demonstrate that as the [*K*^+^]*_P_* concentration gradually increases from the normal physiological concentration ([*K*^+^]*_P_* =6mM), vasodilation ([*K*^+^]*_P_* =11,16mM), inhibited vasodilation ([*K*^+^]*_P_* =21mM), and vasoconstriction ([*K*^+^]*_P_* =26mM) can be achieved, as compared to Fig 2(a).

Finally, we will evaluate the validity of our model through quantification. Experimental results in Fig 5(a), originated from Fig 2(e) in Ref. [15], demonstrate that upon injection of 10mM *K*^+^ concentration into the capillaries, the diameter of the WT blood vessel expands from 15.92*µ*m to 23.51*µ*m (extracted by WebPlotDigitizer). However, the EC Kir2.1 -/- (EC-specific Kir2.1-knockout) only shows minimal fluctuation.

**Fig 5.**
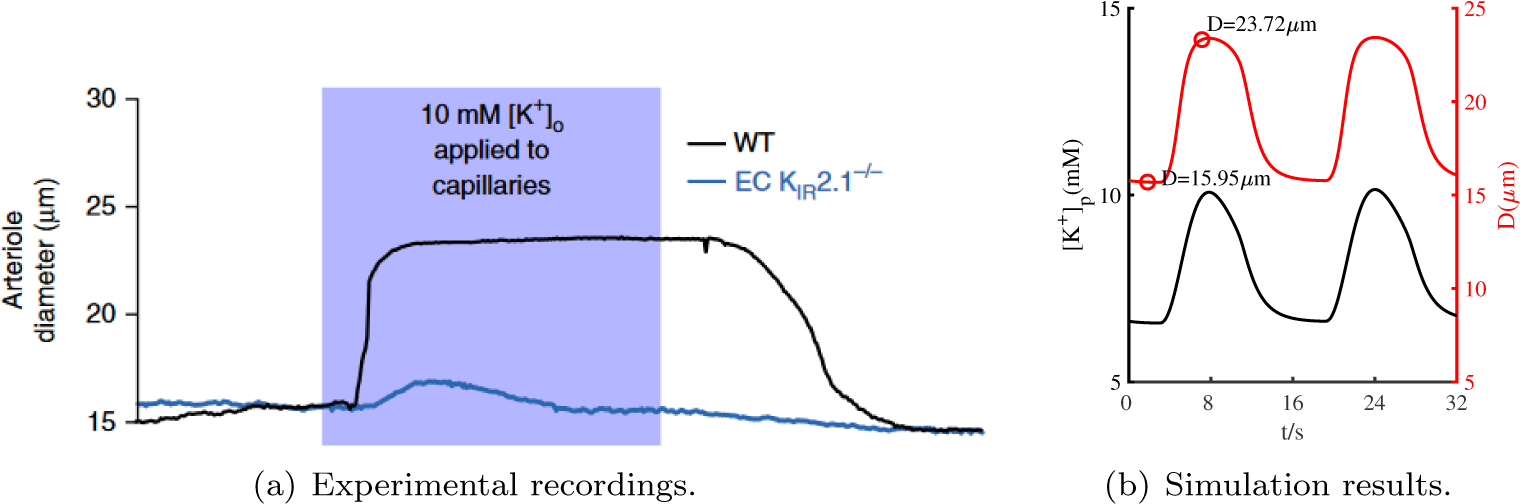
(a) Fig 2(e) in Ref. [15] shows parenchymal arteriole dilation in response to capillary stimulation with 10mM *K*^+^ for 18s in preparations from WT (black trace) and EC KIR2.1-/- (blue trace) mice. (b) Time histories of perivascular potassium ion concentration ([*K*^+^]*_P_*) and vessel diameter (*D*) simulation.

By reducing the value of *k_EET_* in Eqn.(21) (see S1 Appendix), we are able to increase the production of EETs, resulting in faster [*K*^+^]*_P_* generation. This allowed our model to simulate the experimental conditions ranging from [*K*^+^]*_P_* =6mM to 10mM (upper trace in Fig 5(b)), which can be used to derive vascular diameter changes in response (lower trace in Fig 5(b)). Comparison of the simulation results with the experimental data represented by the black line in Fig 5(a), reveals that our model can accurately reproduce the experimental results.

Furthermore, it is essential to present the corresponding simulation results when Kir2.1 channels is present or not and the experimental results from Ref. [15] should also be included for comparison purposes. All the results are summarized in the Fig 6 below. It is evident from the figure that the simulation results accurately reproduce the experimental data, yielding highly consistent results.

**Fig 6.**
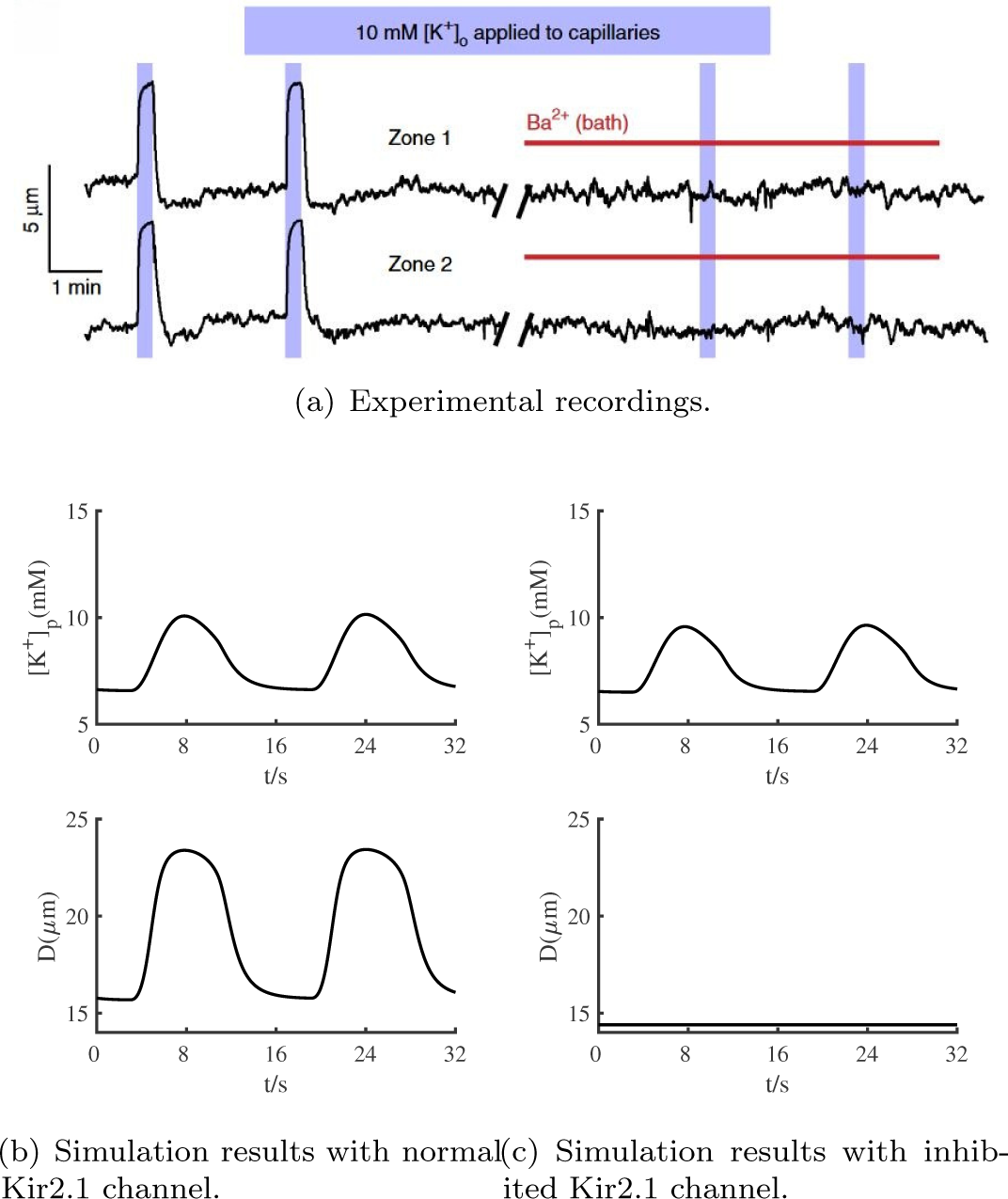
(a) Fig 2(d) in Ref. [15] shows that application of 10 mM *K*^+^ (5 psi) to capillaries produced rapid arteriolar dilation, which was blocked by 30*µ*M *Ba*^2+^. (b) and (c) show the simulation results when the Kir channel is present or inhibited, respectively.

In summary, our newly developed Kir2.1 model has demonstrated excellent agreement with experimental results in terms of existence, quantitative, and qualitative aspects. This constitutes an essential step towards modeling neurovascular coupling units.

### Vasomotion induced by febrile status epilepticus(FSE)

In the FSE rat model, Salehi [41] made the observation that the basolateral amygdala (BLA) exhibits a substantial reduction in both vascular connectivity and diameter, as illustrated in Fig 7. A noticeable reduction in vascular density can be observed in FSE mice compared to normal mice, with an average decrease of approximately 95% across the entire brain. In this experiment, the measurement of blood vessel density encompasses both length and diameter. However, for the purpose of this article, we specifically focus on the changes in diameter as a proxy for changes in density. Here, we examined the impact of acute hyperthermia on neuron firing activities using the neurovascular coupling (NVC) model, which takes into account the influence of temperature. Figure 8 displays numerical simulations that demonstrate how an elevated temperature can trigger changes in spontaneous neuron firing patterns even without any external stimulus input. For instance, at lower temperature ranges [21*^◦^C* to 29*^◦^C*] (Fig 8(a)), neurons exhibit a slow periodic bursting firing state. However, at around 37.5*^◦^C* (Fig 8(b)), neurons switch to a regular spiking pattern, consistent with experimental findings [43]. Furthermore, temperatures above 40*^◦^C* can induce spontaneous febrile status epilepticus(FSE) patterns (Fig 8(c)), similar to what has been previously reported in hippocampal pyramidal neurons [44].

**Fig 7.**
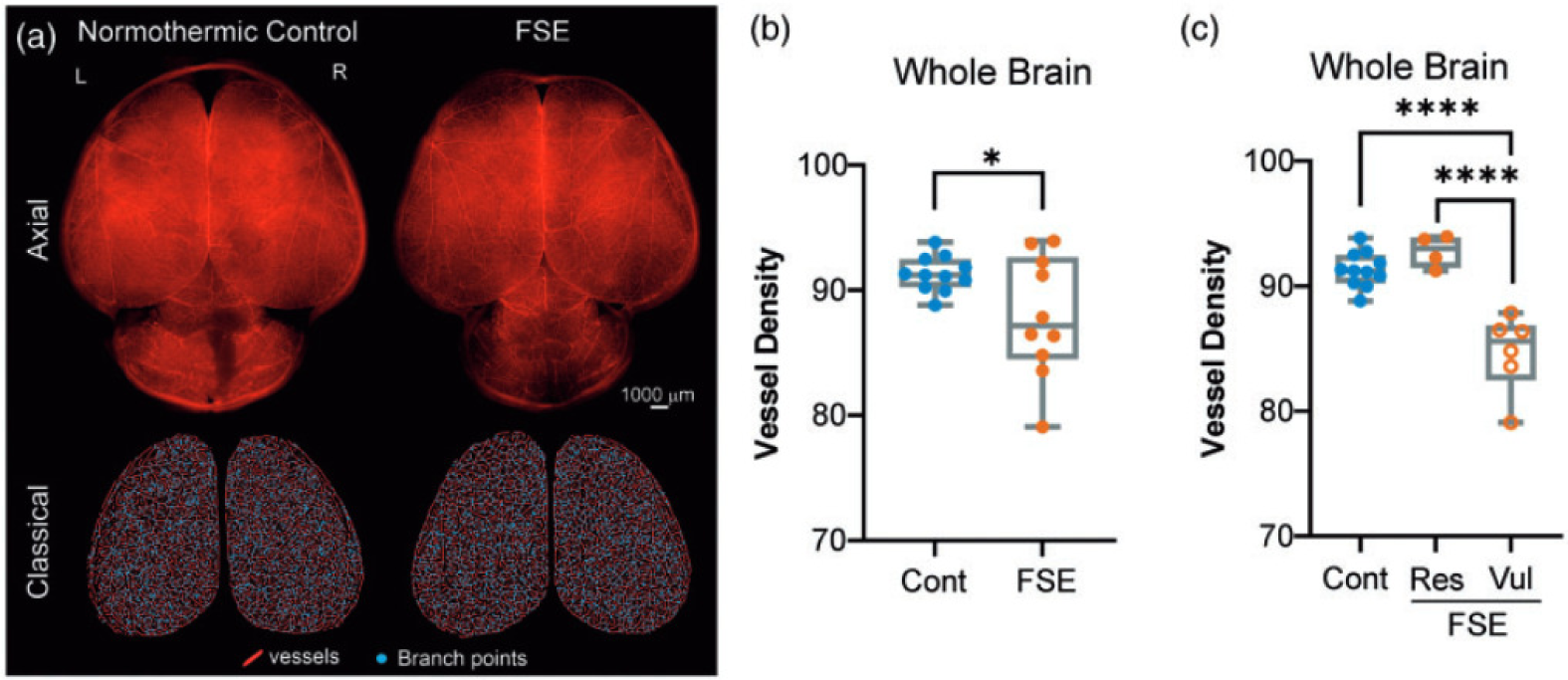
Figures (a), (b), and (c) are adapted from Fig 1 in Ref. [41]. (a) Two hours after the onset of status epilepticus, blood vessel painting was performed on both normothermic control rat pups and those with febrile status epilepticus (FSE). (b) The overall cortical vessel density in the whole brain was found to be significantly different between the two groups. (c) Further analysis, segregating the FSE pups into low or high vessel density groups, revealed significant differences between the vulnerable and resistant vessel density groups, as well as between the controls and the high vessel density group.

**Fig 8.**
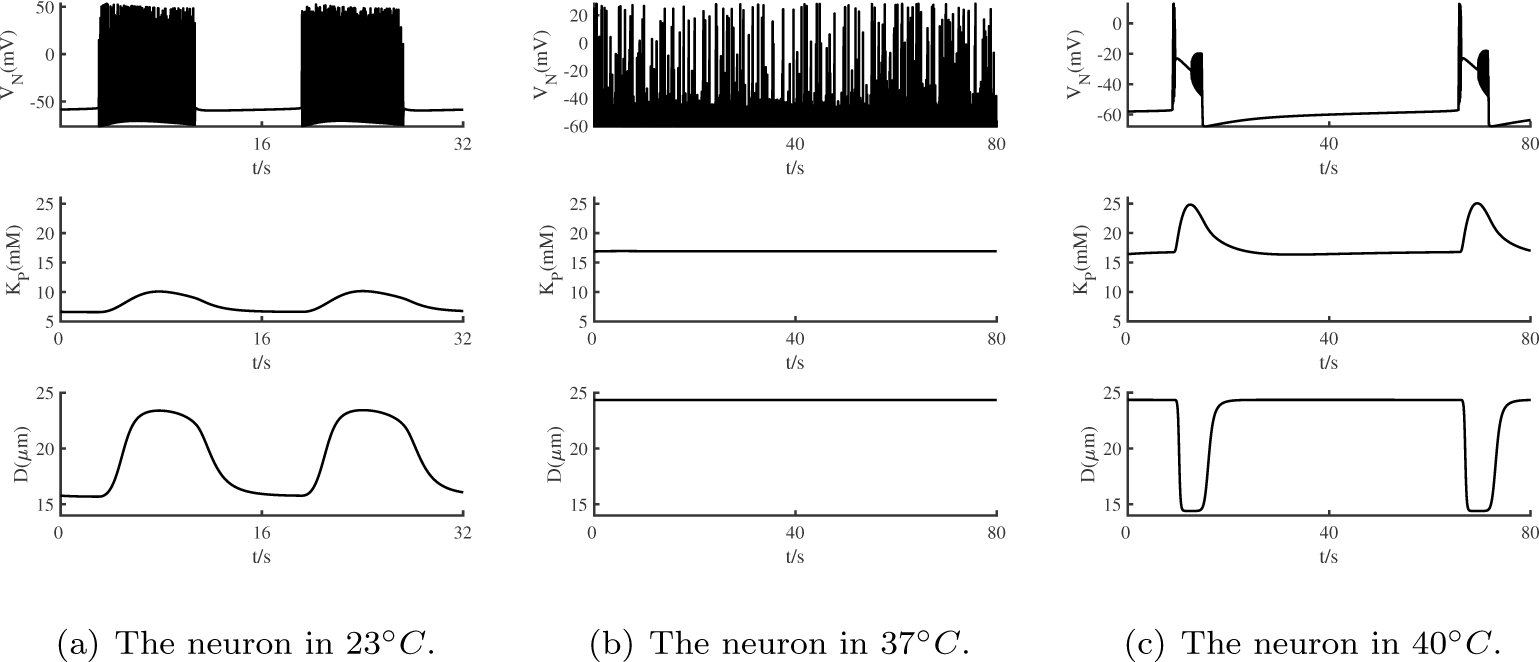
At temperatures of 23 37 and 40 degrees Celsius, the response of the entire neurovascular coupling (NVC) system involves changes in multiple aspects, including the membrane potential of neurons(*V_N_*), the concentration of potassium ions surrounding blood vessels(*K_P_*), and the diameter of the blood vessels(*D*).

Additionally, during the observed depolarization occlusion of neurons in FSE, the diameter of blood vessels decreased to a minimum value of 14.4*µm*, representing 94.7% of the average diameter of blood vessels at 23*^◦^C*. This finding is in accordance with the experimental observations, further validating the accuracy of our study. By incorporating the temperature term into the new NVC model and performing simulations, we have successfully arrived at a conclusion that aligns with the experimental results, thus reaffirming the reliability and validity of the proposed model.

## Conclusion and Discussion

Neurovascular coupling (NVC) is the physiological process by which the brain ensures adequate blood supply to active regions of the brain. It is a complex phenomenon that involves intricate interactions between neurons, astrocytes, pericytes, endothelial cells (ECs), and smooth muscle cells (SMCs). The dynamics of NVC have been studied extensively, but the underlying mechanisms are still not fully understood. In this study, we developed a new model of Kir2.1 channels on endothelial cells, which can sense the effect of capillary *K*^+^-sensing vasomotion and incorporated smooth muscle cells and blood vessel diameter changes into the neuron-glial coupling unit to create a comprehensive neurovascular coupling network. The novel Kir2.1 channel model is rigorously validated from both qualitative and quantitative perspectives and found to be in agreement with experimental results, thus serving as a crucial component in the comprehensive neurovascular coupling network. Moreover, in this model, we incorporate the temperature factor and simulate the vasoconstriction induced by febrile status epilepticus(FSE).

In conclusion, our dynamics model of the neurovascular coupling network provides a valuable tool for studying the complex interactions between neurons, astrocytes, and vascular cells that underlie NVC. By incorporating the effect of capillary *K*^+^-sensing vasomotion, our model sheds light on the role of potassium ions in regulating NVC and provides insights into the mechanisms underlying various brain diseases. Future research can build upon our model to further refine our understanding of NVC and its clinical implications.

Vascular endothelial cells, as a specific type of endothelial cell, are distributed throughout the entire circulatory system, from the heart to the tiniest capillaries. These cells possess distinct functions, including fluid filtration in structures such as the regulation of vascular tension, hemostasis and hormone delivery. In addition to the previously mentioned role of endothelial cells, utilizing their membrane Kir2.1 channel to facilitate vascular movement in response to potassium, there is another mechanism involving the secretion of a substance, nitric oxide (NO), which is also secreted by neurons and contributes to vasomotion. Since NO can traverse neurovascular coupling (NVC) units, it represents a potential direction for future modeling studies.

In this study, the value of *q_t_* in the temperature term is set to 0.2, which differs from the values of 0.1 in Yu’s model [45] and 1 in Du’s model [46]. The reason for this difference is that in this novel NVC model, we aim to replicate the discharge pattern of neurons with depolarization blocking phenomenon under high temperature conditions of 40 degrees Celsius. To achieve this, it was necessary to set *q_t_* at 0.17 or above, based on extensive adjustments and testing. While Yu’s model represents the Hodgkin-Huxley neuron model and Du’s model represents the neuron-astrocyte coupling model, our study has led us to conclude that a value of 0.2 is appropriate for the specific model.

The neurovascular coupling unit is increasingly recognized as a fundamental unit for studying the nervous system and the brain. The aim of this study is to enhance our understanding of the neural control of vascular movement and gain insights into the functioning principles of neurovascular units. To achieve this, we focus on developing a more precise and comprehensive neuron-astrocyte-endothelial-smooth muscle cell coupling model, with the goal of advancing our understanding of the intricate interactions within the neurovascular system.

## Supporting information

**S1 Appendix. Supplementary Materials: Neurovascular Coupling Model Accounting for Temperature.**

## Acknowledgments

We extend our heartfelt gratitude to Professor Yu Yuguo from Fudan University and Wei Yina from Zhejiang Lab for their invaluable guidance and mentorship throughout the course of this research. Their insightful feedback and expertise in modelling significantly shaped our understanding and contributed to the improvement of this work.

